# Texture Profile Analysis and Consumer Sensory Evaluation of Plant-Based and Conventional Breaded Shrimp

**DOI:** 10.64898/2026.07.17.739214

**Authors:** Aeneas O Koosis, Ellen Kuhl

**Author notes:** Corresponding author *Email address:* (Aeneas O Koosis).

## Abstract

Plant-based seafood alternatives could reduce pressure on marine ecosystems, yet texture gaps between plant-based and conventional products limit consumer adoption. This study compared the textural properties and consumer acceptance of plant-based and conventional breaded shrimp to evaluate whether instrumental texture differences predict sensory differences. Texture Profile Analysis (TPA) was performed on four breaded shrimp products (n = 14–15 replicates per product), and consumer sensory evaluation (n = 107) used a within-subjects counterbalanced design assessing hedonic ratings, Just-About-Right attributes, Check-All-That-Apply descriptors, and purchase intent. TPA revealed that plant-based shrimp exhibited significantly higher hardness (14.9 ± 1.0 N vs. 9.6 ± 2.8 N), stiffness (592 ± 40 kPa vs. 382 ± 113 kPa), and chewiness (10.5 ± 0.8 N vs. 2.5 ± 1.1 N) compared to conventional shrimp (all p < 0.0001). However, consumer sensory evaluation revealed no significant differences between products for any hedonic attribute, JAR rating, CATA descriptor, or purchase intent (all p > 0.05 after FDR correction). Despite 1.6-fold greater hardness and 4.2-fold greater chewiness, plant-based breaded shrimp received comparable hedonic ratings to conventional shrimp, suggesting that the breaded format masks mechanical texture differences. Achieving TPA parity may not be necessary for consumer acceptance in breaded seafood applications.

**Highlights:**

- Plant-based shrimp is 1.6× harder and 4.2× chewier than conventional
- Consumers reported no significant sensory differences between products
- Breaded format masks instrumental texture differences
- TPA parity may not be required for consumer acceptance

## 1. Introduction

Global demand for protein is rising, and seafood is positioned to capture a growing share. Per capita fish consumption has nearly doubled since the 1960s, from 10 kg to over 20 kg annually [9], and total seafood demand is projected to reach 186 million tons by 2030, a 20% increase from 2011 levels, driven entirely by aquaculture expansion [13]. Yet supply faces hard limits. Over 35% of global fish stocks are now overfished, and wild capture fisheries have plateaued near 90 million tons for decades [9]. Aquaculture fills the gap but brings its own costs: biodiversity loss, disease outbreaks in densely stocked facilities, and bioaccumulation of heavy metals and microplastics in farmed fish [12]. Against this backdrop, plant-based seafood alternatives have emerged as a potential strategy for meeting protein demand without further straining marine ecosystems.

The sector is growing but remains small. Alternative seafood could supply up to 8% of global seafood demand by 2030 [18], yet consumer adoption lags far behind plant-based beef and chicken. In direct comparisons, plant-based fish analogues score below 40 on 100-point hedonic scales while conventional fish scores 65–75 [2]. Consumers cite off-flavors, unappealing appearance, and, above all, texture: gummy, gelatinous, and lacking the flaky or firm-tender bite of seafood. Replicating seafood texture is technically difficult. Fish and shrimp muscle consists of short, delicate fibers organized into myotomes, a hierarchical fibrous structure that plant proteins do not naturally form [12]. Extrusion, electrospinning, and 3D printing can approximate these structures [6], but results remain inconsistent, and the formulation space is vast [19, 31, 11].

Texture Profile Analysis (TPA) is the standard instrumental method for characterizing food texture. A double-compression test mimics the chewing action of the jaw, yielding parameters– hardness, cohesiveness, springiness, chewiness, resilience–that correlate with sensory texture attributes across many food matrices [5, 26]. For product developers, TPA offers an objective benchmark: match the mechanical profile of conventional shrimp, and consumer acceptance should follow. Recent work on plant-based meat analogues finds that TPA parameters correlate with perceived texture attributes but fail to explain liking [10, 25]. Hardness predicts perceived hardness; it does not predict whether consumers enjoy the product. If the same holds for seafood, optimizing for TPA parity may not be sufficient to ensure consumer acceptance.

Breaded shrimp offers a natural test case. In the US and European markets, breaded and battered formats account for a large share of shrimp consumption; breaded shrimp are convenient, familiar, and may reduce the perceptual salience of product variation [8]. The breading contributes a crispy, crunchy exterior that may mask differences in the protein core. Texture perception follows a temporal sequence during mastication: hardness and crispness dominate early chewing, while chewiness and residual mouthfeel emerge later [17]. If the crispy coating establishes texture expectations before the teeth reach the shrimp, even large mechanical differences in the core may not register with consumers. This raises a fundamental question.

### Can a crispy coating mask substantial textural differences in the protein core without changing consumer perception?

To answer this question, we compared plant-based and conventional breaded shrimp using both TPA and consumer sensory evaluation. Our objectives were to (i) characterize textural differences between products using TPA, (ii) determine whether consumers can distinguish between products using hedonic, Just-About-Right (JAR), and Check-All-That-Apply (CATA) measures, and (iii) evaluate the relationship between instrumental texture measurements and consumer perception. If substantial TPA differences do not translate into sensory differences in breaded formats, achieving instrumental parity may be unnecessary for consumer acceptance.

## 2. Materials and Methods

### 2.1. Samples

Four breaded shrimp products were evaluated in this study (Table 2.1, Figure 1). The primary comparison was between a commercially available plant-based breaded shrimp product (Bayou Best, distributed by Sysco) and a conventional breaded shrimp product (Whole Foods 365, Conv B). Two additional conventional products were included as controls for TPA measurements to assess variability across brands: Gorton’s (Conv A) and SeaPak (Conv C). These were excluded from sensory evaluation due to practical constraints; Conv C showed textural similarity to Conv B, while Conv A broadened the range of conventional products characterized by TPA. All products were purchased frozen and stored at −18°C until testing, with all testing completed within one week of purchase.

**Table 1.** Breaded shrimp products evaluated in this study.

| Product | Analysis | Protein Source | Key Ingredients |
| --- | --- | --- | --- |
| Plant-based | TPA, sensory | Pea protein | Pea protein, calcium lactate, sodium alginate, potato starch |
| Conv A | TPA only | Shrimp | Shrimp, wheat flour, soybean oil |
| Conv B | TPA, sensory | Shrimp | Shrimp, wheat flour, soybean oil |
| Conv C | TPA only | Shrimp | Shrimp, wheat flour, soybean oil |

**Table 2.** Texture Profile Analysis parameters for breaded shrimp products (mean ± SD).

| Parameter | Plant-based | Conv A | Conv B | Conv C |
| --- | --- | --- | --- | --- |
| Stiffness (kPa) | 592.1 ± 39.7<sup>a</sup> | 210.4 ± 59.4<sup>c</sup> | 381.9 ± 112.8<sup>b</sup> | 365.2 ± 112.9<sup>b</sup> |
| Hardness (N) | 14.9 ± 1.0<sup>a</sup> | 5.3 ± 1.5<sup>c</sup> | 9.6 ± 2.8<sup>b</sup> | 9.2 ± 2.8<sup>b</sup> |
| Cohesiveness | 0.79 ± 0.06<sup>a</sup> | 0.53 ± 0.09<sup>bc</sup> | 0.45 ± 0.10<sup>b</sup> | 0.55 ± 0.08<sup>c</sup> |
| Springiness | 0.89 ± 0.04<sup>a</sup> | 0.62 ± 0.04<sup>b</sup> | 0.56 ± 0.15<sup>b</sup> | 0.54 ± 0.15<sup>b</sup> |
| Resilience | 0.74 ± 0.09<sup>a</sup> | 0.53 ± 0.09<sup>bc</sup> | 0.54 ± 0.08<sup>b</sup> | 0.45 ± 0.07<sup>c</sup> |
| Chewiness (N) | 10.5 ± 0.8<sup>a</sup> | 1.8 ± 0.8<sup>b</sup> | 2.5 ± 1.1<sup>b</sup> | 2.6 ± 0.9<sup>b</sup> |

Superscript letters indicate Tukey HSD results. Values sharing the same letter within a row are not significantly different ( $p < 0.05$ ). Conv B (Whole Foods 365) served as the conventional comparator for sensory evaluation.

**Figure 1:**
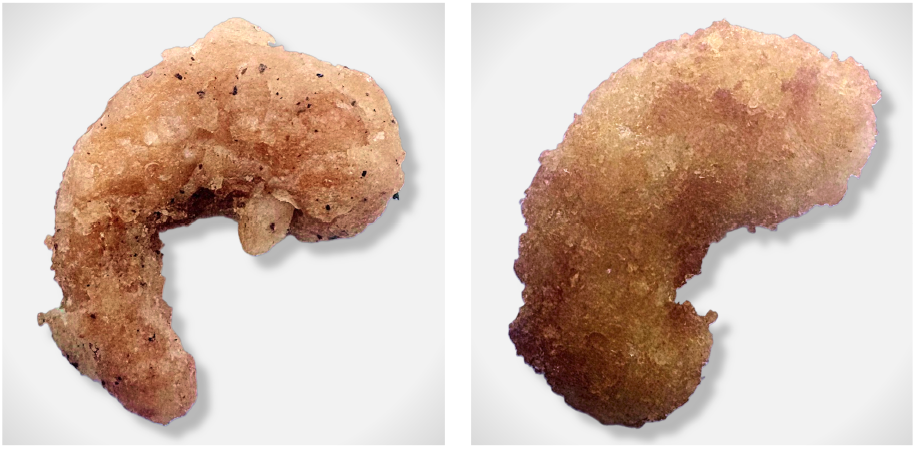
Visual appearance of breaded shrimp products. Plant-based breaded shrimp (left) and conventional breaded shrimp (right) after deep-frying at 177°C. Both products exhibited similar golden-brown coloration and breaded exterior appearance.

### 2.2. Texture Profile Analysis

TPA was performed using a DHR-2 rotational rheometer (TA Instruments) operated in compression mode [7]. The instrument was equipped with an 8-mm diameter cylindrical aluminum probe. Test parameters were pre-test speed 1.0 mm/s, test speed 1.0 mm/s, 50% compression based on individual sample height, trigger force 0.1 N, and 2 s dwell time between compressions. TPA was performed on samples with breading intact to reflect the product as consumed. Testing was conducted within 10 minutes of cooking while samples remained warm. The probe was positioned at the geometric center of each shrimp piece. Fifteen individual shrimp pieces were analyzed per product. Six TPA parameters were calculated from force-time curves [7]:

- *Stiffness* (kPa): Initial slope of force-displacement curve, normalized by probe contact area (50.3 mm^2^)
- *Hardness* (N): Peak force during first compression cycle
- *Cohesiveness* (ratio): Ratio of positive force area during second compression to first compression
- *Springiness* (ratio): Height recovery between compressions
- *Resilience* (ratio): Energy recovered during upstroke relative to energy applied during downstroke
- *Chewiness* (N): Product of hardness × cohesiveness × springiness

Final sample sizes were plant-based (n = 15), Conv A (n = 15), Conv B (n = 14), Conv C (n = 14).

### 2.3. Sensory Evaluation

#### 2.3.1. Participants

Consumer panelists (n = 107) were recruited from Stanford University and the surrounding community via posted flyers and word of mouth. Sample size was determined to achieve 80% power to detect a medium effect size (Cohen’s d = 0.28) with α = 0.05 for paired comparisons. Inclusion criteria required participants to be adults aged 18+ who self-identified as omnivores; omnivore status was verified as a requirement for participation prior to enrollment. Exclusion criteria included allergies to shellfish, soy, or wheat. Stanford University IRB approved this study (eProtocol #87127), exempt under 45 CFR 46.104(d)(6)(i). Participants received a $5 electronic gift card.

#### 2.3.2. Experimental Design

A within-subjects design was employed, with each participant evaluating both plant-based and conventional shrimp. Presentation order was counterbalanced using a counterbalanced AB/BA design [30]. Samples were identified using unique 3-digit codes to blind participants to product identity. Products were deep-fried in vegetable oil at 177°C (350°F) in a commercial kitchen fryer, following manufacturer instructions. For sensory evaluation, samples were served warm within 10 minutes of cooking. Each participant received two pieces of each shrimp. Participants were instructed to cleanse their palate with water and crackers provided between samples, with a one-to-two minute inter-sample interval. No warm-up sample was provided.

#### 2.3.3. Sensory Protocol

For each sample, evaluation proceeded in fixed order [16]: (1) hedonic ratings for appearance, flavor, texture, and overall liking (9-point scale: 1 = dislike extremely, 9 = like extremely); (2) Just-About-Right (JAR) ratings for shrimp flavor intensity, firmness, and chewiness (5-point scale: 1 = much too weak/soft, 3 = just about right, 5 = much too strong/firm) [22]; (3) Check-All-That-Apply (CATA) descriptors [4] (15 sensory attributes: shrimp-like, fishy, salty, savory, bland, firm, tender, chewy, rubbery, juicy, natural-looking, appealing color, fresh aroma, off-flavor, off-odor); and (4) purchase intent (5-point scale: 1 = definitely would not buy, 5 = definitely would buy). CATA attributes were presented in randomized order within each participant to minimize primacy bias.

### 2.4. Statistical Analysis

TPA data were analyzed using one-way ANOVA with Tukey HSD post-hoc tests for multi-product comparisons across all four products. For the primary plant-based vs. Conv B comparison, Welch’s t-test was used to account for unequal variances. Normality of sensory hedonic difference scores was assessed using Shapiro-Wilk tests. As three of four hedonic variables violated normality assumptions (p < 0.05), all sensory comparisons were conducted using non-parametric tests: Wilcoxon signed-rank tests for hedonic ratings, JAR ratings, and purchase intent. Effect sizes were calculated using Cohen’s d for paired samples. CATA frequencies were compared using McNemar’s test with continuity correction.

To control for multiple comparisons across 23 statistical tests (4 hedonic + 1 purchase + 3 JAR + 15 CATA), the Benjamini-Hochberg false discovery rate (FDR) correction was applied at α = 0.05. JAR penalty analysis was conducted to identify deviations from *just about right* that significantly impacted overall liking [21]. Seafood consumption frequency was examined as a potential covariate using linear mixed-effects models. All analyses were performed in Python using scipy, statsmodels, and pingouin libraries. To provide positive evidence of equivalence–rather than merely failing to detect a difference– equivalence testing was conducted using the two one-sided tests (TOST) procedure [15]. Equivalence bounds were set at Cohen’s *d*_*z*_ = ±0.3, corresponding to a small to medium effect size and representing the smallest effect of practical interest for consumer sensory perception. For each paired sensory measure, two one-sided *t*-tests evaluated whether the mean difference was significantly greater than the lower bound and significantly less than the upper bound. Equivalence was concluded when both tests were significant at α = 0.05.

## 3. Results

### 3.1. Texture Profile Analysis

TPA results for all four products are presented in Table 3.1 and Figure 2. Plant-based shrimp exhibited substantially higher values across all six TPA parameters compared to all conventional products. The primary comparison between plant-based and Conv B shrimp revealed highly significant differences across all TPA parameters (p < 0.0001, Welch’s t-test). Plant-based shrimp demonstrated 1.6-fold greater hardness (14.9 vs. 9.6 N), 1.6-fold greater stiffness (592.1 vs. 381.9 kPa), and 4.2-fold greater chewiness (10.5 vs. 2.5 N).

**Table 3.**
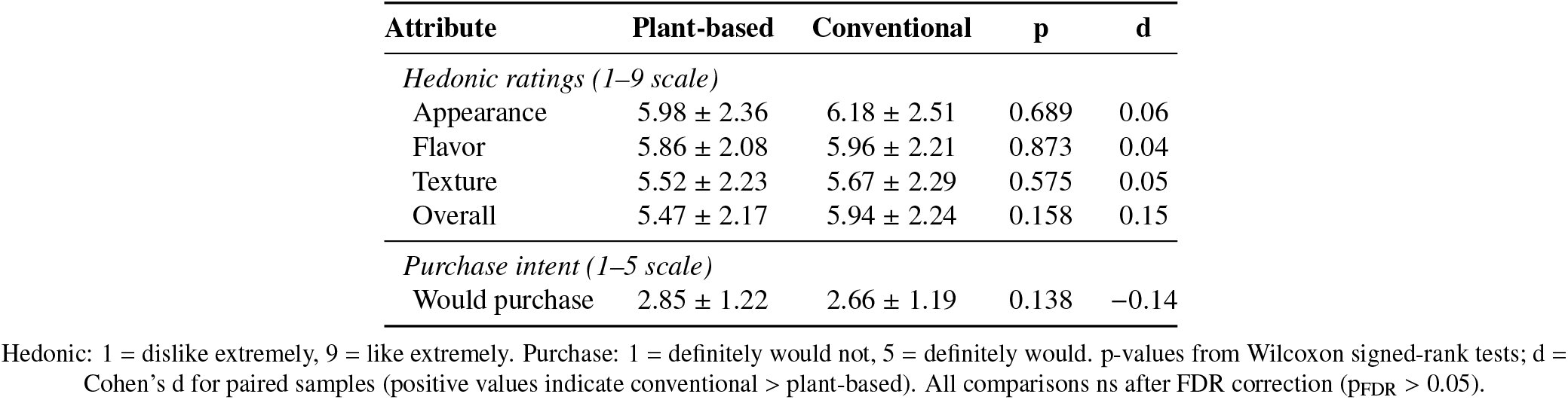
Sensory evaluation results (mean ± SD).

| Attribute | Plant-based | Conventional | p | d |
| --- | --- | --- | --- | --- |
| *Hedonic ratings (1–9 scale)* |  |  |  |  |
| Appearance | 5.98 ± 2.36 | 6.18 ± 2.51 | 0.689 | 0.06 |
| Flavor | 5.86 ± 2.08 | 5.96 ± 2.21 | 0.873 | 0.04 |
| Texture | 5.52 ± 2.23 | 5.67 ± 2.29 | 0.575 | 0.05 |
| Overall | 5.47 ± 2.17 | 5.94 ± 2.24 | 0.158 | 0.15 |
| *Purchase intent (1–5 scale)* |  |  |  |  |
| Would purchase | 2.85 ± 1.22 | 2.66 ± 1.19 | 0.138 | –0.14 |

Hedonic: 1 = dislike extremely, 9 = like extremely. Purchase: 1 = definitely would not, 5 = definitely would. p-values from Wilcoxon signed-rank tests; d = Cohen’s d for paired samples (positive values indicate conventional > plant-based). All comparisons ns after FDR correction ( $p_{FDR} > 0.05$ ).

**Figure 2:**
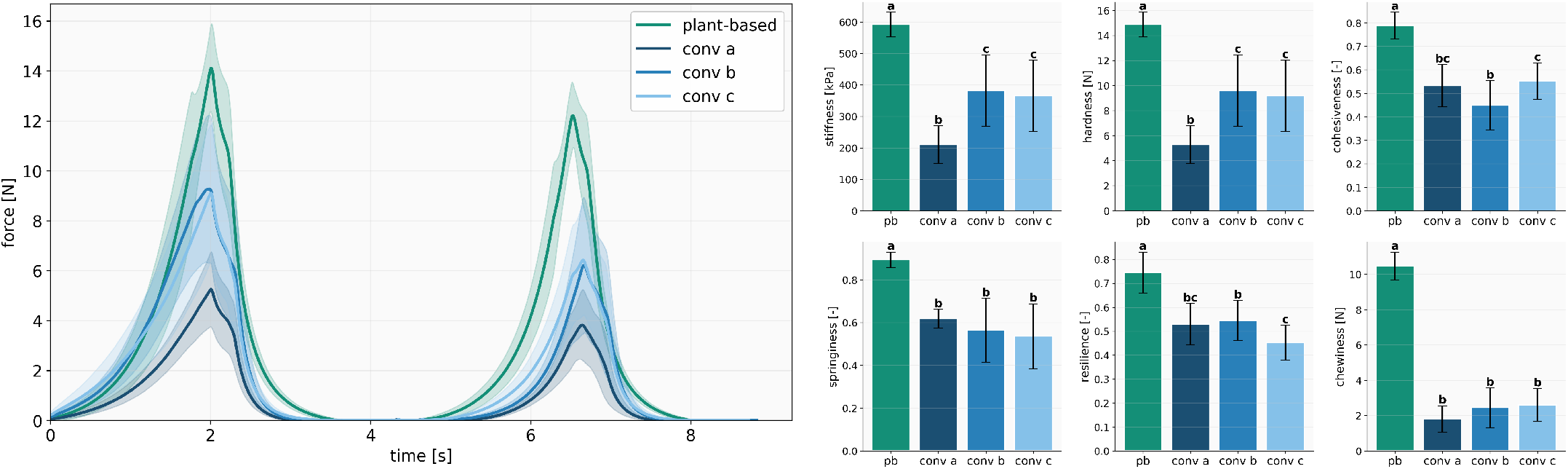
Texture profile analysis. (Left) Mean force-time curves from two-cycle TPA compression tests for four shrimp products. Shaded regions represent standard deviation. (Right) Comparison of six TPA parameters across products. Error bars represent standard deviation. Letters indicate significant differences (Tukey HSD, p < 0.05).

### 3.2. Consumer Sensory Evaluation

#### 3.2.1. Participant Demographics

A total of 107 participants completed the sensory evaluation (47.7% female, 47.7% male, 3.7% non-binary). The sample was predominantly young adults (74.8% aged 18–34). The majority (75.7%) consumed seafood at least 1–2 times per month. Participants represented diverse plant-based consumption habits, ranging from never (8.4%) to daily (3.7%), with the largest group consuming plant-based foods 1–2 times per month (29.9%).

#### 3.2.2. Hedonic Ratings and Purchase Intent

No significant differences were observed between conventional and plant-based shrimp for any hedonic attribute after FDR correction (Table 3.2.2). Both products received moderate ratings (5.47–6.18 on the 9-point scale). Effect sizes were negligible to small (Cohen’s d = 0.04–0.15). Seafood consumption frequency did not significantly influence hedonic ratings when examined as a covariate (all p > 0.05). Purchase intent was numerically higher for plant-based shrimp (2.85 vs. 2.66), though this difference was not statistically significant (p = 0.138). Equivalence testing using the TOST procedure [15] provided positive evidence that the two products were statistically equivalent on four of five sensory measures. With equivalence bounds set at Cohen’s *d*_*z*_ = ±0.3, appearance liking (*d*_*z*_ = 0.05, *p*_TOST_ = 0.007), flavor liking (*d*_*z*_ = 0.04, *p*_TOST_ = 0.004), texture liking (*d*_*z*_ = 0.06, *p*_TOST_ = 0.007), and purchase intent (*d*_*z*_ = −0.14, *p*_TOST_ = 0.047) all demonstrated equivalence. Overall liking did not meet the equivalence criterion (*d*_*z*_ = 0.16, *p*_TOST_ = 0.082), though the observed effect remained small.

Within-subject analysis of hedonic rating differences revealed centered distributions for all attributes (Figure 3). The majority of participants rated both products similarly, with difference scores clustering around zero. For flavor, texture, and overall liking, no systematic preference emerged in either direction (all Wilcoxon p > 0.05).

**Figure 3:**
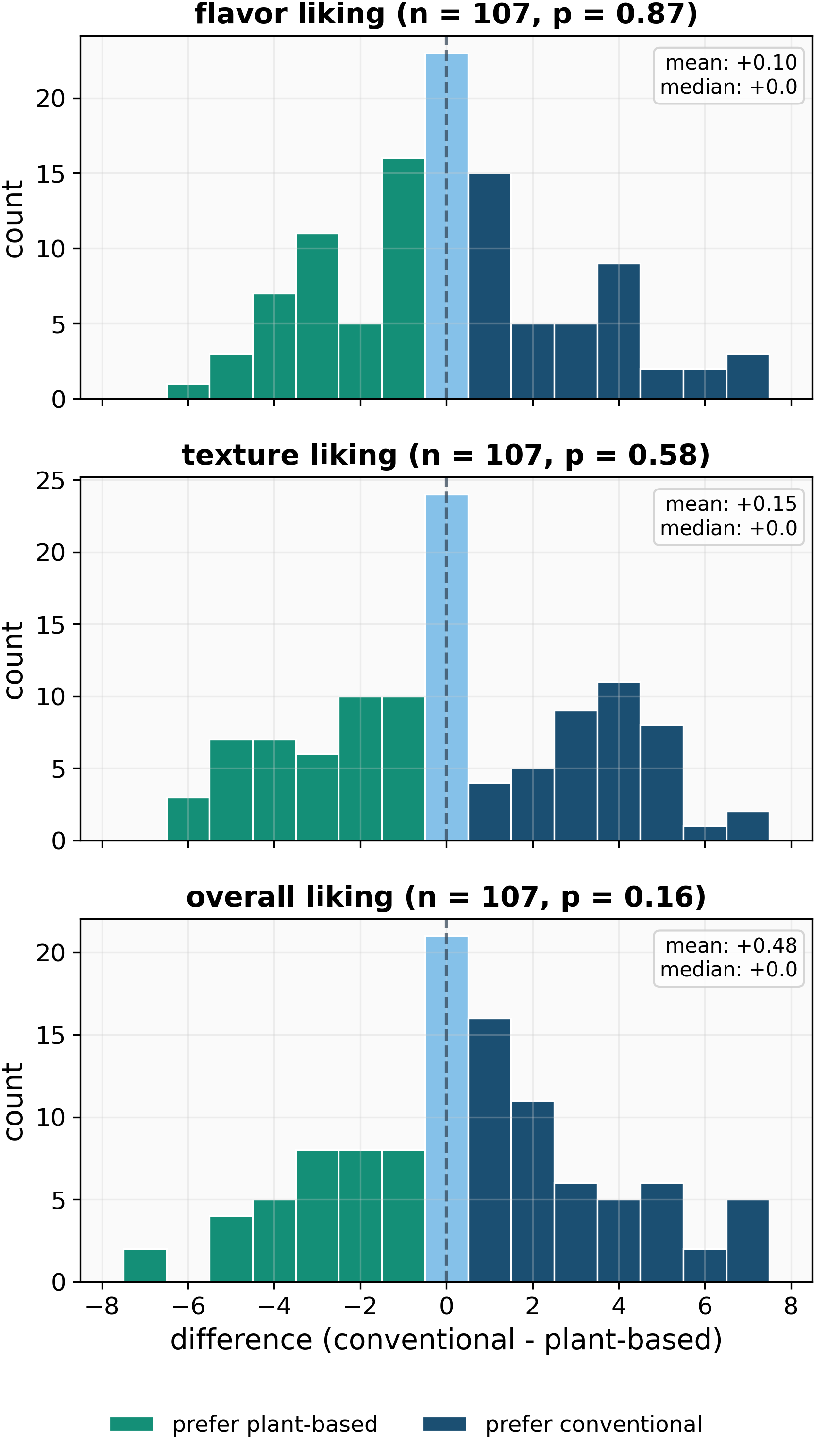
Within-subject hedonic differences. Distribution of paired rating differences (conventional plant-based) for flavor, texture, and overall liking. Blue bars indicate preference for conventional; green bars indicate preference for plant-based. Vertical dashed line marks zero difference. p-values from Wilcoxon signed-rank tests.

#### 3.2.3. Just-About-Right Analysis

No significant differences were observed between products for any JAR attribute (shrimp intensity: p = 0.082; firmness: p = 0.382; chewiness: p = 0.180). For conventional shrimp, the proportion rating attributes as *just about right* was 58.5% (shrimp intensity), 57.5% (firmness), and 57.5% (chewiness). Plant-based shrimp showed similar JAR proportions: 57.0% (shrimp intensity), 43.9% (firmness), and 50.0% (chewiness) (Figure 4).

**Figure 4:**
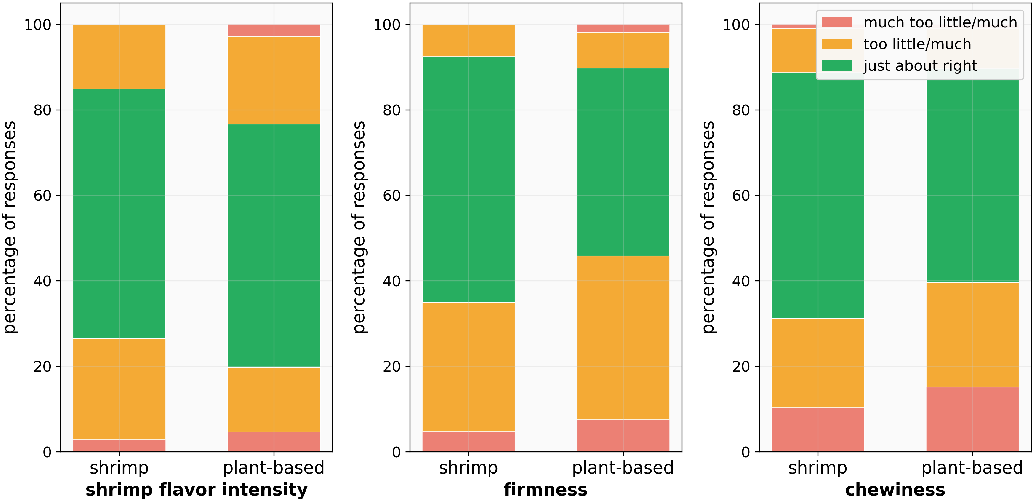
Just-About-Right distributions. JAR ratings for shrimp flavor intensity, firmness, and chewiness. Green = just about right; yellow = slight deviation; red = strong deviation from optimal.

Penalty analysis revealed that deviations from JAR significantly impacted overall liking for both products. For conventional shrimp, *too soft* (penalty = 1.01, p = 0.033) and *not chewy enough* (penalty = 1.20, p = 0.015) decreased overall liking. For plant-based shrimp, firmness deviations in both directions reduced overall liking with *too soft* (penalty = 1.13, p = 0.008) and *too firm* (penalty = 2.42, p = 0.001), as did insufficient chewiness (penalty = 0.91, p = 0.041).

#### 3.2.4. CATA Analysis

Check-All-That-Apply analysis revealed no significant differences between products for any of the 15 sensory attributes after FDR correction (all p_FDR_ > 0.05, McNemar’s test). The most frequently selected attributes for conventional shrimp were bland (45.8%), salty (44.9%), and off-odor (39.3%). For plant-based shrimp, the most frequent were natural-looking (43.9%), tender (43.0%), and salty (36.4%). The largest difference between products was for chewy, which was selected more frequently for plant-based (28.0%) than conventional shrimp (16.8%), though this difference did not reach significance (p_raw_ = 0.074, p_FDR_ = 0.533). Correspondence analysis (Figure 5) visualized attribute associations for each product; however, no individual attribute differed significantly between products after correction.

**Figure 5:**
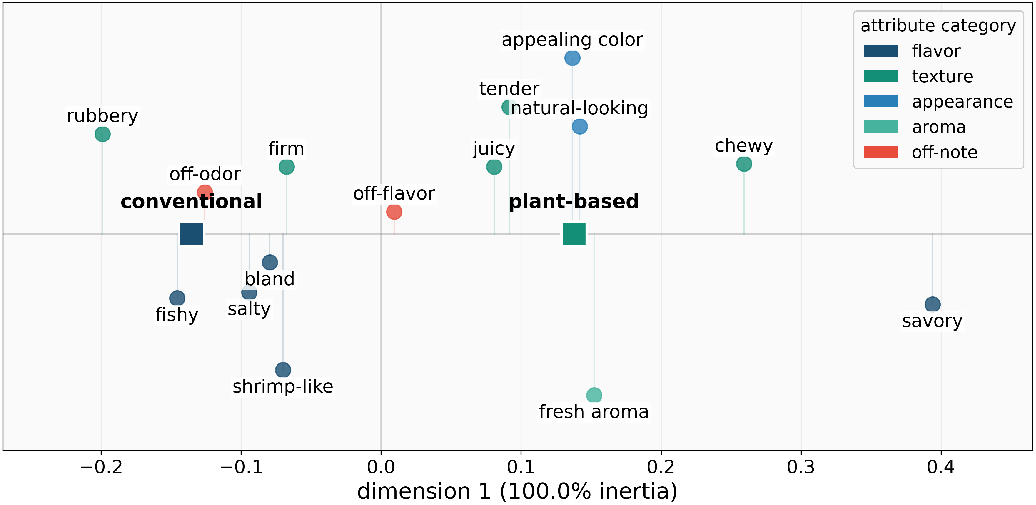
Correspondence analysis. One-dimensional correspondence analysis map of CATA sensory attributes. Because two products yield one non-zero correspondence dimension, proximity should be interpreted along the horizontal axis.

### 3.3. Instrumental-Sensory Comparison

While TPA revealed highly significant differences across all texture parameters (p < 0.0001), consumers did not distinguish between products on any sensory measure, including texture liking (p = 0.575), firmness JAR (p = 0.382), and chewiness JAR (p = 0.180). The instrumental differences (1.6–4.2-fold) appear to fall within the range of acceptable variation for breaded shrimp, or the breading masks underlying textural differences during mastication.

## 4. Discussion

The central finding is an instrumental-sensory mismatch: *Instrumental measurements detected large textural differences between plant-based and conventional shrimp, but consumers did not report statistically significant differences on any sensory measures*. Plant-based shrimp was 1.6× harder, 1.8× more cohesive, and 4.2× chewier, differences that would likely matter more in an unbreaded product. This result fits a broader pattern. TPA parameters often correlate with perceived texture attributes–consumers who rate a product as “hard” tend to be eating something with high instrumental hardness–but this correlation does not always extend to liking. Across 27 commercial meat analogues, TPA hardness, chewiness, and cohesiveness tracked sensory ratings of those same attributes, but none explained overall liking; liking depended on cross-modal factors like meaty flavor, juiciness, and fattiness [10]. A similar disconnect emerged for meatballs: animal-based samples were significantly harder, more cohesive, and chewier by TPA, yet consumers perceived no texture differences between categories [25]. Hybrid plant-meat burgers with significantly different TPA profiles received identical liking scores [20]. Mechanically matched burgers received divergent meatiness and overall liking ratings [14].

Texture perception during mastication follows a temporal sequence: hardness, crackliness, and crispness dominate at the start of chewing, while attributes like stickiness and residual chewiness emerge later [17]. When biting into breaded shrimp, the first sensation is fracture of the fried coating. This crisp exterior may set texture expectations before the teeth reach the protein core. Deep-fried breaded coatings show strong correlations between coating structure and perceived crispness [29]. This suggests that the coating, rather than the filling, may drive initial texture perception. The breading also creates structural heterogeneity: a crispy exterior over a softer interior. In heterogeneous foods, mechanical contrast between components shapes overall perception in ways that single-phase TPA cannot capture [23]. TPA compresses the whole sample, whereas oral processing does not necessarily experience the sample as a homogeneous mass.

For plant-based seafood development, this result is encouraging. Texture remains a primary barrier to consumer acceptance [1], and replicating the fibrous structure of fish muscle is technically difficult [19, 31]. Recent work on breaded plant-based fish filets found that analogues scored lower than animal-based samples, with pasty and gummy texture negatively impacting overall liking [3]. We observed no such penalty. Breaded plant-based shrimp received comparable hedonic and texture ratings to conventional shrimp. This shifts the practical target: Instead of matching the mechanical profile of conventional shrimp, which current formulations clearly do not, the goal may be to achieve acceptable texture within a breaded format. Our data suggest this plant-based product approaches that target, though the marginal difference in overall liking indicates room for improvement. Purchase intent was numerically higher for the plant-based product (2.85 vs. 2.66), though not significantly. The key result is that the plant-based product was not penalized despite being mechanically stiffer and chewier.

This does not diminish the value of TPA. For uncoated products, including sashimi analogues, shrimp salads, and unbreaded fish fillets, mechanical texture likely plays a larger role in perception, and TPA may better predict consumer response [7]. Nor does it mean texture does not matter. Penalty analysis showed that deviations from *just about right* firmness and chewiness reduced liking for both products. Recent work on AI-generated burgers found that texture independently drives liking even when flavor and appearance are controlled [27]. Texture may function as a threshold rather than a continuous predictor: once texture is acceptable, other factors such as flavor, appearance, and juiciness may matter more to overall liking. *The lesson is not that instrumental methods fail, but that they measure the product, not the eating experience*. Researchers developing breaded plant-based seafood should treat TPA as a screening tool, rather than the sole optimization target. Open-source benchmarking protocols [28] and validation against human sensory data are necessary steps.

Several limitations apply. We tested one plant-based product; other formulations may perform differently. The sample skewed young, with 75% under 35, and was drawn from a university community. Participants were blinded to product identity; informed conditions shift preferences toward or against plant-based options depending on consumer values [24]. Finally, we tested only a breaded format. The instrumental-sensory disconnect we observed may not hold for unbreaded preparations.

## 5. Conclusion

Breaded plant-based shrimp was 1.6× harder and 4.2× chewier than conventional shrimp by TPA, yet consumers did not rate the products differently on any sensory measure. This instrumental-sensory disconnect has practical implications. For plant-based seafood developers targeting breaded formats, acceptable consumer texture may not require full mechanical parity with conventional shrimp. The plant-based formulation achieved comparable hedonic ratings for appearance, flavor, and texture, but a small overall liking gap persisted. These findings suggest that achieving full TPA parity may not be necessary for consumer acceptance in breaded formats, although generalization to other products requires further testing. For researchers, the results highlight a boundary condition for TPA. The method reliably detects mechanical differences, but those differences do not always reach consumer perception, particularly when coatings, batters, or heterogeneous structures intervene between instrument and mouth. TPA remains useful for screening and quality control, but sensory validation is essential before treating instrumental targets as product specifications. Whether this disconnect extends beyond breaded formats remains an open question. For uncoated applications like shrimp salads, ceviche, or sashimi-style products, texture likely plays a more direct role in perception, and TPA may better predict consumer response. Future work should compare breaded and unbreaded plant-based shrimp head-to-head, and investigate the specific mechanisms by which coating structures could be masking core texture differences during mastication.

## Data availability

De-identified sensory data, TPA results, and analysis scripts are available at https://github.com/LivingMatterLab/AI4Food.

## Acknowledgments

This research was conducted under Stanford University IRB Protocol #87127. It was supported by the Stanford Bio-X Postdoctoral Research Grant, the Stanford Bio-X Snack Grant, the Stanford SDSS Accelerator Grant, the NSF CMMI Award 2320933, and the ERC Advanced Grant 101141626.

